# C3: An R package for cross-species compendium-based cell-type identification

**DOI:** 10.1101/267880

**Authors:** Md Humayun Kabir, Djordje Djordjevic, Michael D. O’Connor, Joshua W. K. Ho

## Abstract

Cell type identification from an unknown sample can often be done by comparing its gene expression profile against a gene expression database containing profiles of a large number of cell-types. This type of compendium-based cell-type identification strategy is particularly successful for human and mouse samples because a large volume of data exists for these organisms. However, such rich data repositories often do not exist for most non-model organisms. This makes transcriptome-based sample classification in these species challenging. We propose to overcome this challenge by performing a *cross-species* compendium comparison. The key is to utilise a recently published cross-species gene set analysis (XGSA) framework to correct for biases that may arise due to potentially complex homologous gene mapping between two species. The framework is implemented as an open source R package called C3. We have evaluated the performance of C3 using a variety of public data in NCBI Gene Expression Omnibus. We also compared the functionality and performance of C3 against some similar gene expression profile matching tools. Our evaluation shows that C3 is a simple and effective method for cell type identification. C3 is available at https://github.com/VCCRI/C3.

## Introduction

The key question we seek to address in this article is *how can we identify the cell-type of a biological sample given its gene expression profile*? This question commonly arises when investigating a novel cell population resulting from differentiation of pluripotent stem cells or isolation of a cell population in a non-model organism. The most popular bioinformatics approach is a compendium-based identification approach, in which the unknown sample’s gene expression profile is used as a query profile against a large gene expression compendium consisting of many cell types. A number of tools have been developed to perform such a task, such as GEMINI [1], ProfileChaser [2], ExpressionBlast [3] and CellMortage [4]. All these tools work in a similar fashion: match the query gene expression profile or a gene set against a database of gene expression profiles to identify its best matches. Importantly, most of these tools implicitly assume there is a one-to-one correspondence between genes in the query sample and the compendium sample, which can be violated when comparing data from different species. Beyond supporting filtering for genes with one-to-one homology mapping across species, none of the current tools effectively handle a cross-species query in a statistically rigorous fashion.

Therefore, when using currently available tools it is important to always use a database of the same species as the query sample. This is often practically impossible because most publicly available data sets are only available for a small number of species. Let’s take as an example one of the largest public gene expression repositories, the NCBI Gene Expression Omnibus (GEO) [5]. As of March 2017, there were more than 57,000 GEO series (GSE) generated by microarrays or RNA-Seq. Collectively, these data are a valuable resource for researchers to discover new biological insights. Nonetheless, most of these GSE data sets were generated from just two species: *Homo sapiens* (human) and *Mus musculus* (mouse). In fact, around two thirds of these GSE data sets are derived from human or mouse samples (Figure 1). The other third come from more than 1,300 species, with only 33 species having over 100 GSE (Figure 1). In other words, while it is possible to curate a useful gene expression compendium for human and mouse, it is practically impossible for other species, especially non-model organisms.

**Figure 1.**
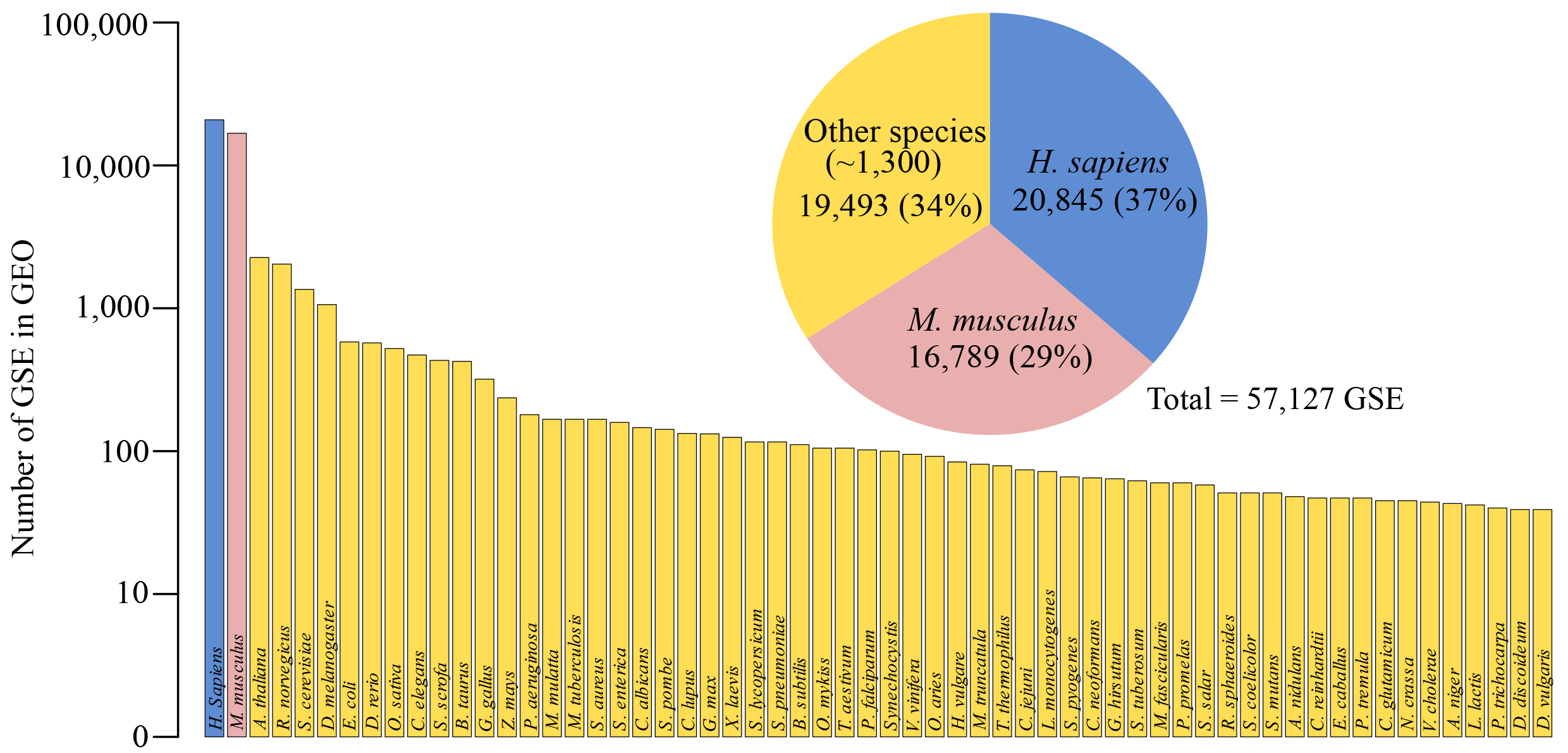
Summary of GSE based on species in NCBI GEO. The pie chart shows the total number of GSE for *H. sapiens* (blue), *M. musculus* (pink) and all other species (orange). The bar plot shows the top 60 species according to the number of GSE in NCBI GEO.

We propose to alleviate this lack of species-specific compendia by performing a *cross-species* cell identification, where a query profile is matched against a database of samples which come from different organisms. A key challenge to implementing such a cross-species analysis scheme is that many pairs of species, especially those that are evolutionary distant, can have complex “many-to-many” homologous gene relationships. Failure to properly account for the homology gene mapping can lead to statistical biases [6].

In this article, we present a new open source R package – C3 – that implements this cross-species compendium-based cell type identification approach using a recently developed cross-species gene set analysis method called XGSA [6]. XGSA has been shown to reduce the false positive bias while still maintain good statistical power for gene sets affected by highly complex homology structures. Using C3, we can harness the large collection of human and mouse public data as a resource to identify unknown cell types for a wide variety of species. We demonstrate the effectiveness of C3 using a large collection of GEO data. We also compare its performance with other similar tools.

## Methods

### C3: a new R package for cross-species cell-type identification

C3 is an open source R package for identifying an unknown cell-type from its gene expression profile based on a large compendium of gene expression data that can be derived from different species. A key aspect of this approach is that it is most useful when the compendium represents many different tissue or cell types, preferably from a well-studied organism such as human or mouse. Examples of public data sources that can be used to form this kind of compendium include ENCODE [7, 8] and GTEx [9]. The full description of the method implemented in C3 is described in detail in the rest of this section, but an overview of the framework can be found in Figure 2. Briefly, C3 first identifies genes considered to be specifically-expressed genes in the query and the compendium profiles, by removing genes ubiquitously expressed across these expression profiles. Next, C3 performs XGSA between the query gene set and each of the compendium gene sets to account for “many-to-many” gene relationships, and thereby determine which compendium gene sets are statistically enriched in the query gene set. A *p*-value is reported for each compendium sample. The cell-types of the most highly ranked compendium gene sets (according to *p*-value) are then used to predict the cell-type of the query profile. C3 is available at https://github.com/VCCRI/C3.

**Figure 2.**
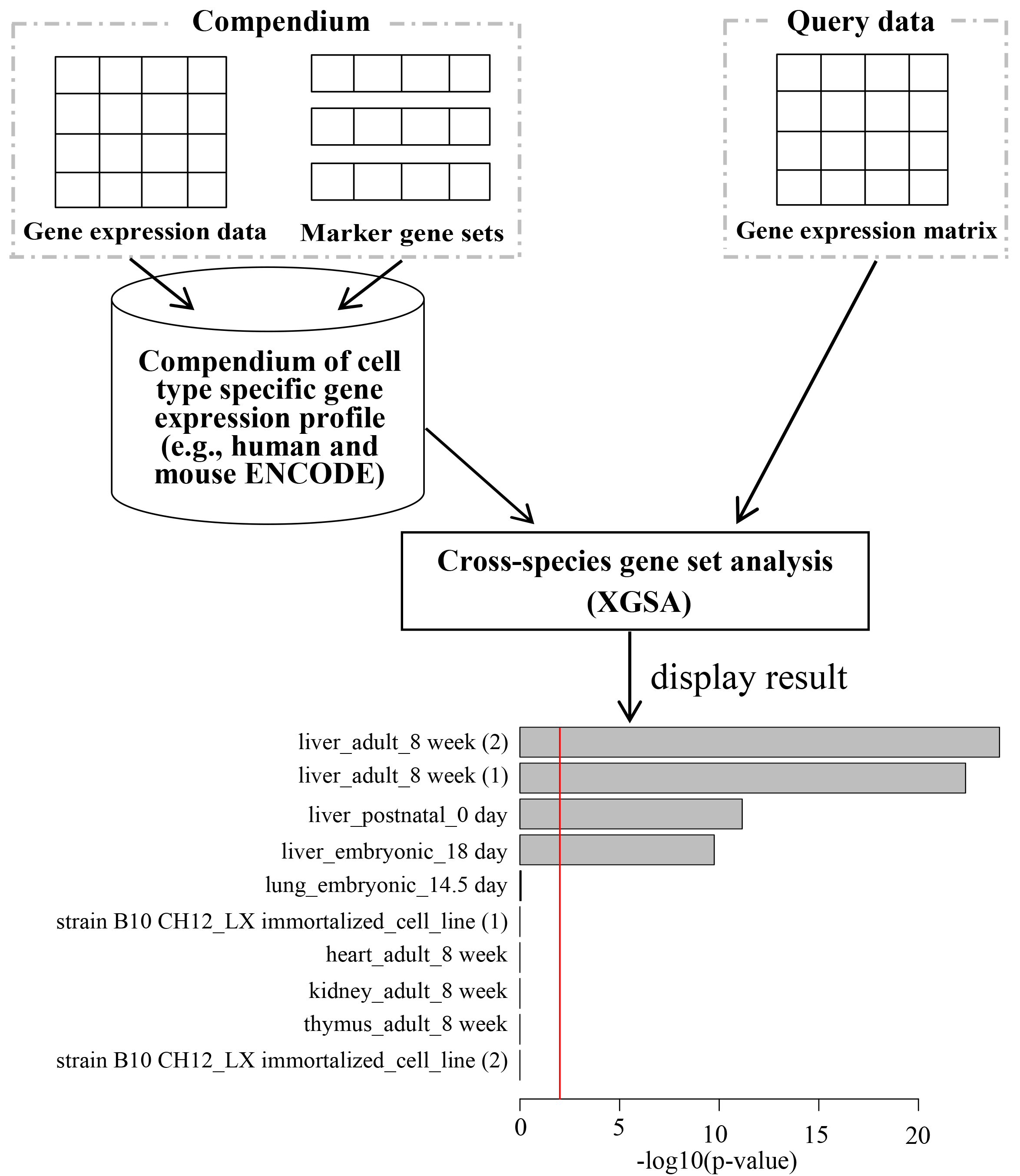
Overall workflow diagram of C3.

### The human and mouse gene expression compendia

For both mouse and human, we constructed a large compendium of tissue-specific genes using RNA data from the ENCODE project. ENCODE gene expression data, summarised as FPKMs, were obtained for human (hg19; 144 tissues or cell lines) [7] and for mouse (mm9; 94 tissues or cell types) [8]. Most tissues or cell types in the ENCODE data set are represented by more than one replicate. We combined replicates of the same tissue or cell type by calculating the mean expression value for each gene. If a compendium is constructed from multiple data sources, we only consider genes that are common among all data sets.

### Identification of specifically expressed genes in the query and compendium data

Using the compendium data, for each sample in the compendium we identified sets of highly-expressed genes that are specific to each sample using two parameters: *n* – the number of highly expressed genes to consider for marker gene status; *t* – the proportion of samples a marker gene can appear in before it is discarded as non-unique/non-specific. Using these two parameters we could identify then remove genes that are consistently highly expressed (within the top *n* highly expressed genes in each sample) in more than *t* × 100% of samples. The goal of this step is to remove ubiquitously expressed genes such as housekeeping genes. The remaining gene sets should be enriched for cell-type specific genes. To identify the highly-expressed specific genes within the query data set, first we identified the top *n* highly expressed genes. We then removed the ubiquitously expressed genes identified by the compendium from the top *n* expressed genes. When the query sample species is different from the species used to create the compendium, we use XGSA to identify the homologs of the set of ubiquitously expressed genes for the query cell species. We then remove this set of gene homologs from the query cell top expressed genes.

### XGSA

To provide the required input for XGSA, all genes names are first converted to ENSEMBL gene IDs. XGSA then applies a simple statistical method that computes a conservative *p*-value based on Fisher’s Exact test. This approach takes into account the homology gene mapping structure between two cross-species gene sets [6]. If the two compared gene sets are from the same gene sets, the resulting *p*-value is identical to that of a standard gene set test based on a Fisher’s Exact test. The package then performs Benjamini-Hochberg multiple testing corrections on the raw *p*-values, and reports and visualises the -log10 of the corrected *p*-values.

### Comparison with ExpressionBlast

For the comparison with ExpressionBlast, we used brain, kidney and liver sample data sets from the *R. norvegicus* species [10]. We identified the specific highly expressed genes for each of the sample tissue types using our C3 package by setting parameter values as *n* = 1000 and *t* = 0.10. Among these specific highly expressed genes, we have selected the top 100 expressed genes based on their expression values. We used this set of highly expressed tissue specific genes with *log2* expression values as the input to the ExpressionBlast web tool. In this way we have tested each of the three tissue types against both the human and mouse organism using ExpressionBlast.

## Results

### Evaluation of C3

To evaluate the performance of C3, we collected gene expression profiles from four GEO data series (GSE43013 [10], GSE74754 [11], GSE78770 [12], and GSE53393 [13]), which collectively contain data from 13 different species (*B. taurus*, *C. familiaris*, *C. porcellus*, *E. caballus*, *E. europaeus*, *F. catus*, *M. musculus*, *O. cuniculus*, *R. norvegicus*, *S. scrofa*, *D. rerio*, *T. truncates*, and *M. mulatta*) across five different tissue types (brain, kidney, liver, blood, and skeletal muscle). We tested whether C3 could correctly identify the cell type of the samples when compared against a human compendium or a mouse compendium constructed from ENCODE data [7, 8]. For comprehensiveness, we tested two combinations of parameters in C3 (*n* and *t*). The summary result is shown in Figure 3 and the detailed results are shown in the Supplementary materials [see Supplementary Tables 1-2]. Overall, baring a few exceptions which will be discussed below, C3 was able to consistently identify the correct or the most closely related cell type across all species (Figure 3).

**Figure 3.**
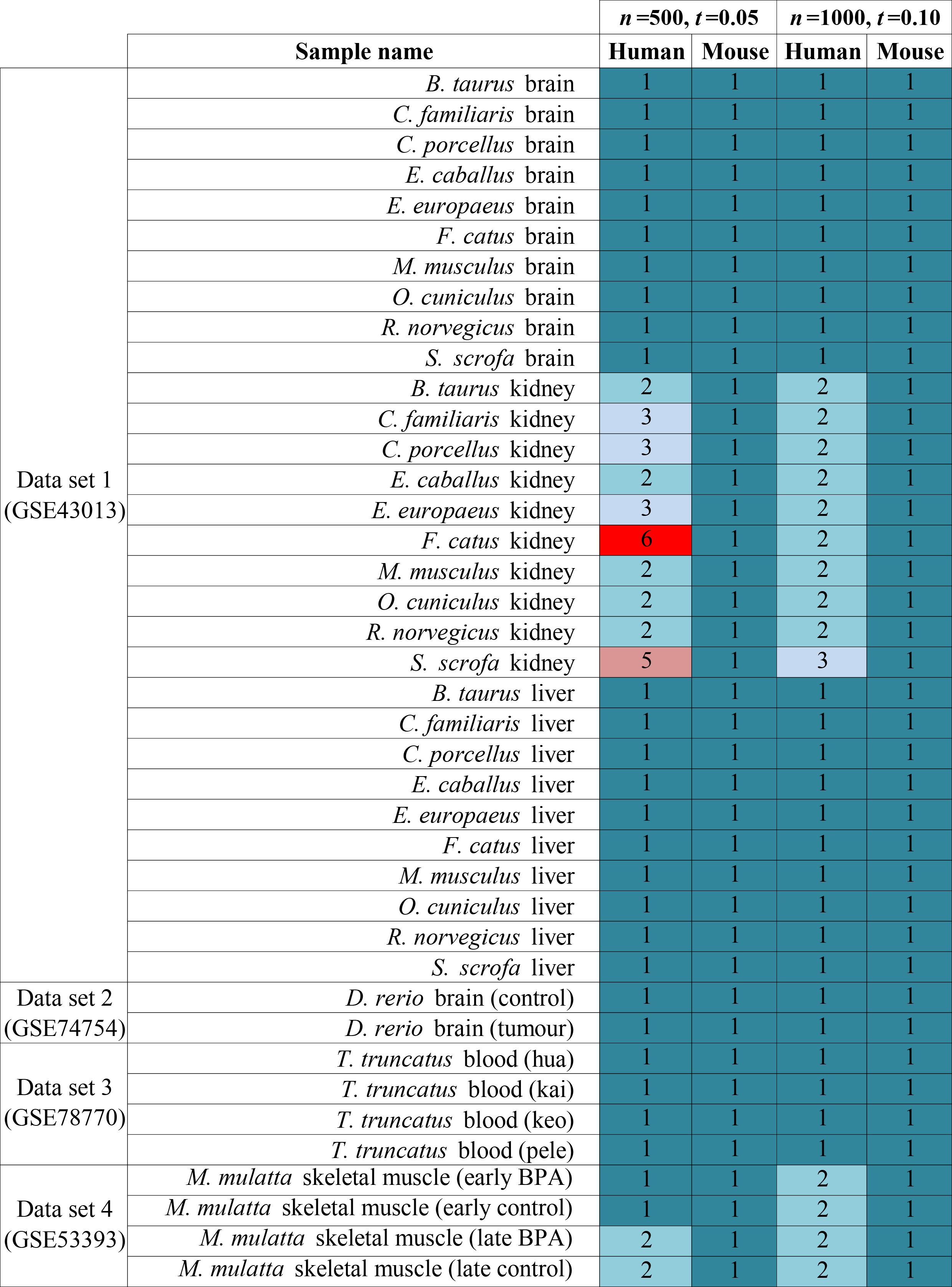
Evaluation of C3. Gene expression profiles of tissues from 13 different organisms were selected from four GEO data sets. These profiles were used to evaluate whether C3 could correctly identify its cell type of the sample when compared against a human ENCODE compendium (Human) or a mouse ENCODE compendium (Mouse). *n*: top number of highly expressed genes; *t*: cut-off threshold value; 1 = Statistically significant and in top position; 2 = Statistically significant but in top 2-3rd position; 3 = Statistically significant but in top 4-5th position; 4 = Not statistically significant but in top position; 5 = Not statistically significant but in top 2-5th position; 6 = Not statistically significant and not in 2-5th position

GSE43013 [10] contains a gene expression data set from three different tissue types (brain, kidney and liver) in 33 mammalian species, among which 10 have homology mapping information available via ENSEMBL. C3 could correctly identify the cell types in all the brain and liver samples across all 10 species. For the kidney data, C3 correctly identified the cell type when compared against the mouse compendium across 10 species, but was much less effective when compared against the human compendium. Interestingly, this comparison against the human compendium resulted in most of the kidney gene sets being identified as liver samples ahead of the human kidney samples. As both of these tissues are highly vascularised, it may be that gene expression profiles from blood and blood vessel cells within the kidney samples confound the analysis against the human compendium.

We also tested three more GSE datasets that contained data from 3 additional species; *D. rerio* (GSE74754; brain) [11], *T. truncates* (GSE78770; blood) [12], and *M. mulatta* (GSE53393; skeletal muscle) [13]. Through these analysis C3 correctly identified the cell types of *D. rerio* brain and *T. truncates* blood. The *M. mulatta* skeletal muscle samples were correctly identified by C3 when they compared to the mouse compendium but were not as effectively identified using the human compendium (top hit was heart/tongue sample) (Figure 3). As with the kidney, skeletal muscle is also highly vascularised – and this could be the cause of the misidentification of the *M. mulatta* skeletal muscle sample using the human compendium.

Overall, a total of 160 C3 analyses were performed (80 against the mouse compendium and 80 against the human compendium) using two combinations of *n/t* parameters (i.e., 500/0.05 and 1000/0.1). Notably, all the cell type identity predictions made by C3 using the mouse compendium were correct for at least one of the parameter combinations (i.e., typically at least 1000/0.1 if not also 500/0.05). For comparison against the human compendium: correct predictions were made for 67.5% of the queries, and for a further 25% of the queries the correct prediction was ranked second or third by C3 (i.e., the correct prediction was in the top 3 positions 92.5% of the time using the human compendium). Only 1 out of the 80 predictions made by C3 using the human compendium (0.625%; *F. cattus*, *kidney*) did not include the correct identification in the top 5 predictions. Notably, only two cell types were not predicted correctly (i.e., as the top prediction): kidney and skeletal muscle. These tissues are both highly vascularised, and this may be a confounding factor when comparing against human samples. However, as shown in Figure 3, all the kidney and skeletal muscle datasets were correctly identified when compared against the mouse compendium.

### Comparison with other similar software programs

A comparison of the features of C3 and other similar methods is illustrated in Table 1. The four similar methods discussed are primarily web-based with only GEMINI offering a Python command-line version. GEMINI lacks the ability to perform cross-species cell type identification. It uses level 3 gene expression datasets from The Cancer Genome Atlas (TCGA) project [14]. CellMontage can compare only the expression data from similar microarray platforms. As a result, neither of these methods could be included in our comparative analysis. ProfileChaser supports cross-species analyses using NCBI HomoloGene for only 6 species, and uses only the set of genes that have one-to-one human homology mapping. However, ProfileChaser searches only the curated GEO DataSets (GDS) (support only 1,815 GDS) for similar biological conditions based on differential gene expression from reduced set of gene expression features. We were unable to meaningful include this tool in our comparative analysis.

**Table 1.** Comparison of software features of C3 and other similar methods.

|  | C3 | ExpressionBlast | ProfileChaser | GEMINI | CellMontage |
| --- | --- | --- | --- | --- | --- |
| **Cross-species method** | Ensembl BioMart portal, complete homology structure using XGSA | Inparanoid, handles multiple orthologues using closest value of input gene | One-to-one human homolog | Not supported | Not mentioned |
| **How many species** | As many as ENSEMBL mapping | 8 | 6 | - | - |
| **Input** | Gene expression matrix | Max 100 differentially expressed genes with expression values | Gene expression matrix | Gene expression matrix | Gene expression matrix with raw expression values |
| **User interface** | R command line | Web | Web | Web and Python command-line | Web |
| **Availability** | Open source | Free | Free | Free | Free |
| **Application** | General | General | Specific to GDS | Level 3 gene expression from TCGA project | Specific to similar microarray platforms |
| **Dependency** | Previously made compendium | Differentially expressed genes | Reduced set of gene expression features | Reduced dimension of expression profile | UniGene names for gene ids |

The only C3 alternative we are aware of that can compare a transcriptomic profile to a compendium of data across species in order to identify an unknown cell type is ExpressionBlast [3]. ExpressionBlast is a web-based tool that takes a maximum of one hundred differentially expressed genes with their expression values, and compares it to microarray data from 8 different species on GEO. For cross-species comparisons, ExpressionBlast uses homologous gene groups from InParanoid and handles multiple homologs using the closest expression value of the input gene. In contrast, C3 is an open source R package that takes gene expression profiles as input. C3 leverages XGSA to perform cross-species analysis between any of species in the growing list of species in Ensembl Compara (currently 93 species).

To compare the performance of ExpressionBlast with C3, we analysed the brain, kidney and liver sample data from *R. norvegicus* (GSE43013) [10] using both methods, as the rat is one of the eight species supported by ExpressionBlast. For C3, we tested against the human and mouse compendiums with parameter values *n*=1000 and *t*=0.10. For ExpressionBlast, we inputted the100 highly expressed tissue specific genes with their *log2(FPKM+1)* expression values. The summary results for C3 and ExpressionBlast are shown in Table 2, and the detailed results are presented in Supplementary Table 2 (for C3) and Supplementary Figure 1 (for ExpressionBlast). From the comparative test results, it is clear that C3 can identify cell type at least as accurately as ExpressionBlast. Nonetheless, C3 is has markedly greater flexibility than ExpressionBlast in that it can handle the whole query gene expression profile, it can be applied to data from a wide range of organisms, and its R package enables it to be easily incorporated into any analytical pipeline.

**Table 2.**
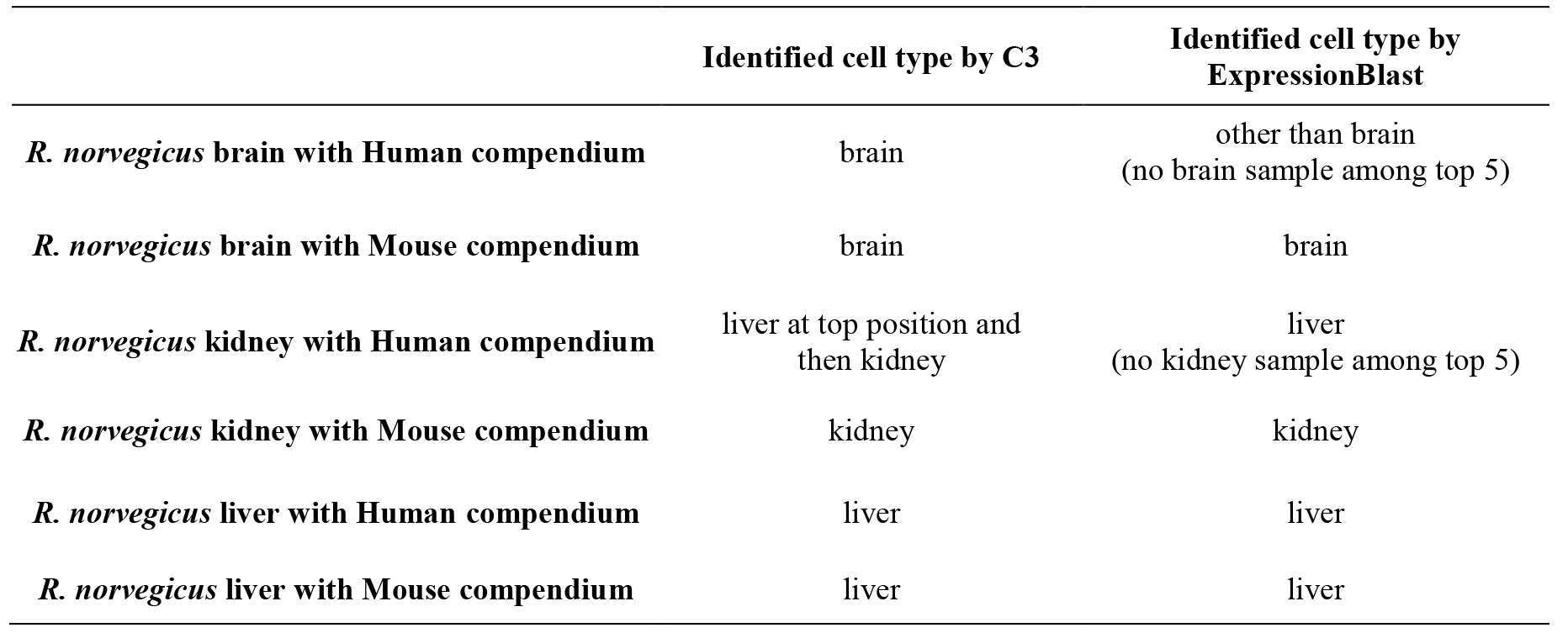
Comparison of cross-species cell type identification using C3 and ExpressionBlast.

|  | Identified cell type by C3 | Identified cell type by ExpressionBlast |
| --- | --- | --- |
| *R. norvegicus* brain with Human compendium | brain | other than brain<br>(no brain sample among top 5) |
| *R. norvegicus* brain with Mouse compendium | brain | brain |
| *R. norvegicus* kidney with Human compendium | liver at top position and then kidney | liver<br>(no kidney sample among top 5) |
| *R. norvegicus* kidney with Mouse compendium | kidney | kidney |
| *R. norvegicus* liver with Human compendium | liver | liver |
| *R. norvegicus* liver with Mouse compendium | liver | liver |

## Discussion

This work highlights the utility of cross-species analysis in cell-type identification using a gene expression compendium-based approach. This is particularly important when considering that the majority (two thirds) of transcriptomic data in the GEO database is from human and mouse, with the remaining third of data shared between over 1,000 organisms (Figure 1), most of which have very scant genomic resources. Our aim with C3 was to leverage the many published data sets from the well characterised human and mouse organisms to identify an unknown cell type from a potentially poorly characterised organism.

Recently we have used this approach to identify that a novel PAX7+ cell population in lizard *Anole carolinensis* is highly similar to muscle satellite cells in human and mouse [15]. As another real-life application, we have recently used the C3 approach to demonstrate that a ROR1+ cell population derived from human pluripotent stem cells is similar to lens epithelial cells in both human and mouse [16]. Both examples highlight the power of C3 in determining or confirming the identity of a cell type using a compendium of gene expression profiles from different species.

C3 can only correctly identify the cell type of an unknown transcriptomic profile if a similar cell type is represented in the compendium. With this in mind, the quality, variety and size of the compendium is paramount and future work should investigate larger compendiums such as based on ARCHS4 [17], as well as domain specific compendiums such as for identifying cancer subtypes.

## Conclusion

Overall, we demonstrated that C3 can prioritise identification of the correct corresponding cell type as the most significant hit. We believe C3 should facilitate rapid cell type identification for less characterised species, or for poorly characterised cell types obtained from stem cell differentiation strategies.

## Authors’ contributions

J.W.K.H. initiated the project; M.H.K. designed the method, implemented the package, performed evaluation and wrote the manuscript; D.D. contributed to method design and software testing; M.D.O’C and J.W.K.H. supervised the whole project and revised the manuscript. All authors read and approved the final manuscript.

## Acknowledgements

M.H.K. is supported by a UWS Postgraduate Research Award (International). J.W.K.H is supported by a Career Development Fellowship by the National Health and Medical Research Council (1105271) and a Future Leader Fellowship by the National Heart Foundation of Australia (100848).

## Supplementary material

**Supplementary Table 1** Detail test result with different species’ different cells/tissues with *n*=500, *t*=0.05

**Supplementary Table 2** Detail test result with different species’ different cells/tissues with *n*=1000, *t*=0.10

**Supplementary Figure 1** Test result screenshot of *R.norvegicus* sample datasets using ExpressionBlast: (a) and (b) show results for brain dataset with *H. sapiens* and *M. musculus* respectively; (c) and (d) show results for kidney dataset with *H. sapiens* and *M. musculus* respectively; (e) and (f) show results for liver dataset with *H. sapiens* and *M. musculus* respectively.

